# Examination of gene loss in the DNA mismatch repair pathway and its mutational consequences in a fungal phylum

**DOI:** 10.1101/2021.04.13.439724

**Authors:** Megan A Phillips, Jacob L Steenwyk, Xing-Xing Shen, Antonis Rokas

**Affiliations:** Department of Biological Sciences, Vanderbilt University, Nashville, TN 37235, USA; Institute of Insect Sciences, Ministry of Agriculture Key Lab of Molecular Biology of Crop Pathogens and Insects, College of Agriculture and Biotechnology, Zhejiang University, Hangzhou 310058, China

**Keywords:** mutation rate, Ascomycota, microsatellite, phylogenetics, powdery mildew, DNA repair

## Abstract

The DNA mismatch repair (MMR) pathway corrects mismatched bases produced during DNA replication and is highly conserved across the tree of life, reflecting its fundamental importance for genome integrity. Loss of function in one or a few MMR genes can lead to increased mutation rates and microsatellite instability, as seen in some human cancers. While loss of MMR genes has been documented in the context of human disease and in hypermutant strains of pathogens, examples of entire species and species lineages that have experienced substantial MMR gene loss are lacking. We examined the genomes of 1,107 species in the fungal phylum Ascomycota for the presence of 52 genes known to be involved in the MMR pathway of fungi. We found that the median ascomycete genome contained 49 / 52 MMR genes. In contrast, four closely related species of obligate plant parasites from the powdery mildew genera *Erysiphe* and *Blumeria*, have lost between 5 and 21 MMR genes, including *MLH3*, *EXO1*, and *DPB11*. The lost genes span MMR functions, include genes that are conserved in all other ascomycetes, and loss of function of any of these genes alone has been previously linked to increased mutation rate. Consistent with the hypothesis that loss of these genes impairs MMR pathway function, we found that powdery mildew genomes with higher levels of MMR gene loss exhibit increased numbers of mononucleotide runs, longer microsatellites, accelerated sequence evolution, elevated mutational bias in the A|T direction, and decreased GC content. These results identify a striking example of macroevolutionary loss of multiple MMR pathway genes in a eukaryotic lineage, even though the mutational outcomes of these losses appear to resemble those associated with detrimental MMR dysfunction in other organisms.

**Significance Statement:** The DNA mismatch repair pathway corrects nucleotide base errors that occur during the replication of DNA; loss of these genes leads to cancer. We examined the conservation of the DNA mismatch repair pathway’s genes across more than 1,000 species in a fungal phylum and found a lineage of powdery mildews, a group of fungi that infect the leaves of plants, which have experienced extensive loss of multiple, otherwise highly conserved, genes. The genomes of these powdery mildews show elevated rates of diverse types of mutation, raising the hypothesis that these organisms have diversified while lacking genes thought to be essential for the accurate replication of DNA.

## Introduction

An ensemble of DNA repair pathways and cell cycle checkpoints is responsible for detecting and repairing DNA damage, ensuring faithful maintenance of the genome (Friedberg et al. 2005; Giglia-Mari et al. 2010). Among DNA repair pathways, the DNA mismatch repair (MMR) pathway is one of the best characterized (Marti et al. 2002). The MMR pathway is responsible for repairing bases that were incorrectly paired during DNA replication via five steps: recognition, incision, removal, re-synthesis, and ligation (Fig. 1A) (Hsieh & Zhang 2017; Fukui 2010; Marti et al. 2002). The MMR pathway is highly conserved in both bacteria and eukaryotes; cells that experience reduction or loss of function in this pathway have increased levels of mutation, as seen in cancer and drug-resistant fungal pathogen strains (Fukui 2010; Campbell et al. 2017; Billmyre et al. 2017, 2020; Dos Reis et al. 2019).

**Fig. 1.**
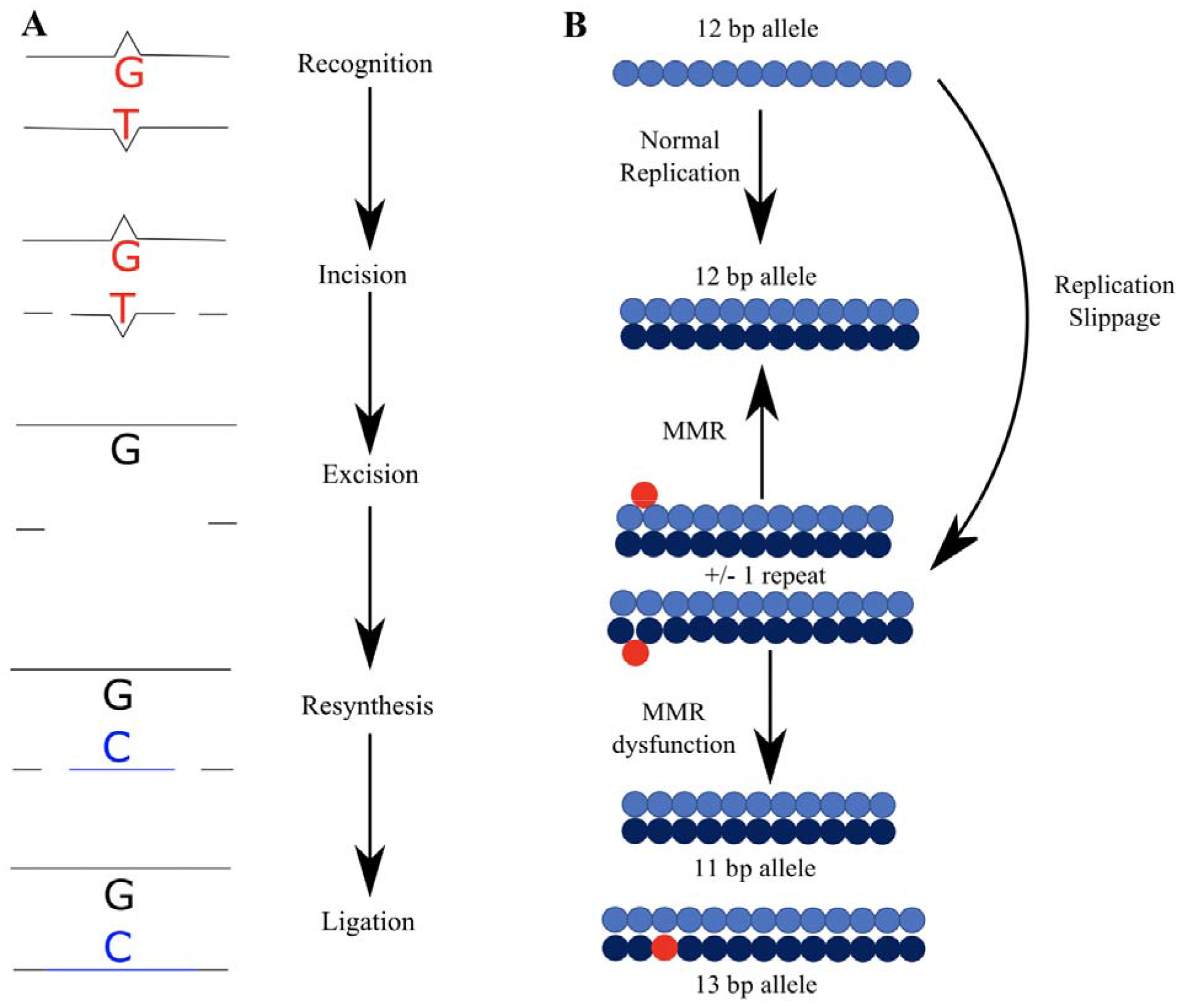
The DNA Mismatch Repair (MMR) pathway corrects mismatched bases produced during DNA replication and prevents instability in microsatellites. (A) The pathway is comprised of five conserved steps: recognition of mispaired bases by a sliding clamp, incision of the DNA strand by an endonuclease, excision of the incorrectly paired bases, resynthesis of the DNA strand, and ligation of the newly synthesized segment to the DNA strand. Table S2 includes a full list of MMR genes and their categorization into one of the five steps. (B) The MMR pathway also recognizes base or repeat addition and skipping errors during replication and corrects them. However, dysfunction in this pathway can leave replication slippage unrepaired, leading to the and the addition or deletion of base pairs or repeats, especially in highly repetitive regions such as microsatellites.

Although DNA repair genes are generally highly conserved, certain fungal lineages have been reported to have a more limited repertoire, particularly within the phylum Ascomycota. For example, budding yeasts (subphylum Saccharomycotina) and fission yeasts (Taphrinomycotina) have fewer DNA repair genes than filamentous fungi (Pezizomycotina) (Milo et al. 2019; Shen et al. 2020). Furthermore, DNA repair genes that were lost from budding yeasts and fission yeasts are more likely to also be lost in filamentous fungi (Milo et al. 2019). One lineage that has experienced extensive losses in its repertoire of DNA repair genes is the *Hanseniaspora* genus of budding yeasts (Steenwyk et al. 2019). *Hanseniaspora* species have undergone punctuated sequence evolution and have accumulated large numbers of diverse types of substitutions, including those associated with specific gene losses such as UV damage, suggesting that DNA repair is impaired by the high levels of DNA repair gene loss. These findings suggest that DNA repair genes are not universally conserved across fungi and that their loss is compatible with long-term evolutionary survival and diversification of fungal lineages.

One well-established consequence of MMR dysfunction is mutation in microsatellite regions of the genome. Microsatellites are repetitive tracts of DNA, with motifs 1-6 bp long repeated at least five times (Beier et al. 2017). Microsatellites are typically highly polymorphic between individuals and are commonly used as markers in population biology, forensics, paternity testing, and tumor characterization (Richman 2015). Due to their repetitive nature, microsatellites are prone to experiencing polymerase slippage, which is usually corrected by the MMR pathway (Fig. 1B) (Ellegren 2004; Richman 2015). If the MMR pathway does not recognize these errors, as is the case in cancer, microsatellite instability (MSI) can occur (Campbell et al. 2017). MSI is defined by a hypermutable phenotype resulting from a loss of function in the MMR pathway (Boland & Goel 2010). Instability in microsatellites trends towards elongation in these regions, but contraction can also occur (Ellegren 2004).

Beyond increased mutation in microsatellite regions, aberrant function of the MMR pathway is associated with genome-wide signatures of genetic instabilities (Boland & Goel 2010; Billmyre et al. 2017, 2020). MMR mutations have been implicated in the development of hypermutant and ultrahypermutant human cancers, which constitute approximately 15% of human tumors and less than 1% of tumors, respectively (Campbell et al. 2017). Interestingly, very few tumors with low mutation rates contained mutations in the MMR pathway, whereas more than a third of hypermutant tumors and virtually all the ultrahypermutants contained mutations in MMR genes (Campbell et al. 2017). Hypermutant tumors had high levels of MSI suggesting their hypermutant phenotype is due, at least in part, to MMR dysfunction (Campbell et al. 2017). Hypermutation has also been observed in fungal pathogen strains that have lost MMR pathway genes, potentially driving within-host adaptation and the evolution of drug resistance. For example, Rhodes et al. (2017) found that hypermutation caused by mutations in three MMR pathway genes, including *MSH2*, resulted in a rapid increase in the mutation rate of the human pathogenic fungus *Cryptococcus neoformans*, contributing to infection relapse. Similarly, Billmyre et al. (2017) sequenced multiple strains of the human pathogenic fungus *Cryptococcus deuterogattii* (phylum Basidiomycota) and found that a group of strains with mutations in the *MSH2* gene experienced higher rates of mutation when compared with closely related strains harboring an intact *MSH2* gene. Hypermutation in *C. deuterogattii* mediated rapid evolution of antifungal drug resistance (Billmyre et al. 2017, 2020).

In contrast to MMR gene loss in the microevolutionary context of genetic or infectious disease, the extent of MMR gene loss across lineages spanning multiple species remains understudied. To determine the macroevolutionary impact of MMR gene conservation and loss, we characterized patterns of MMR gene presence and absence in the fungal phylum Ascomycota (Fig. 2). We found that the MMR pathway was highly conserved across Ascomycota, with the median species having 49 / 52 MMR genes present. However, we found that *Blumeria graminis* and species in the powdery mildew genus *Erysiphe* (subphylum Pezizomycotina, class Leotiomycetes), a group of obligate plant parasites, had many fewer MMR genes and a faster rate of sequence evolution than their relatives and most other fungal taxa. Specifically, *Erysiphe necator* has lost 9 MMR genes, *Erysiphe pisi* has lost 21 MMR genes, *Erysiphe pulchra* has lost 7 MMR genes, and *Blumeria graminis* has lost 5 MMR genes (Fig. 3). In contrast, species closely related to *Erysiphe* and *Blumeria* have lost only 1 – 2 MMR genes, consistent with the high degree of MMR gene conservation in the rest of the phylum. Evolutionary genomic analyses revealed that MMR gene losses in *Erysiphe* and *Blumeria* (hereafter referred to as higher loss taxa or HLT) are associated with a proliferation of mononucleotide runs and elongation of microsatellites of all motif lengths, both of which are hallmarks of MMR pathway dysfunction. Reflecting these losses, *Erysiphe* and *Blumeria* genomes also have more pronounced mutational biases and accelerated rates of mutation. These results suggest that obligate plant parasites in the genera *Blumeria* and *Erysiphe* have diversified while lacking otherwise highly conserved MMR genes.

**Fig. 2.**
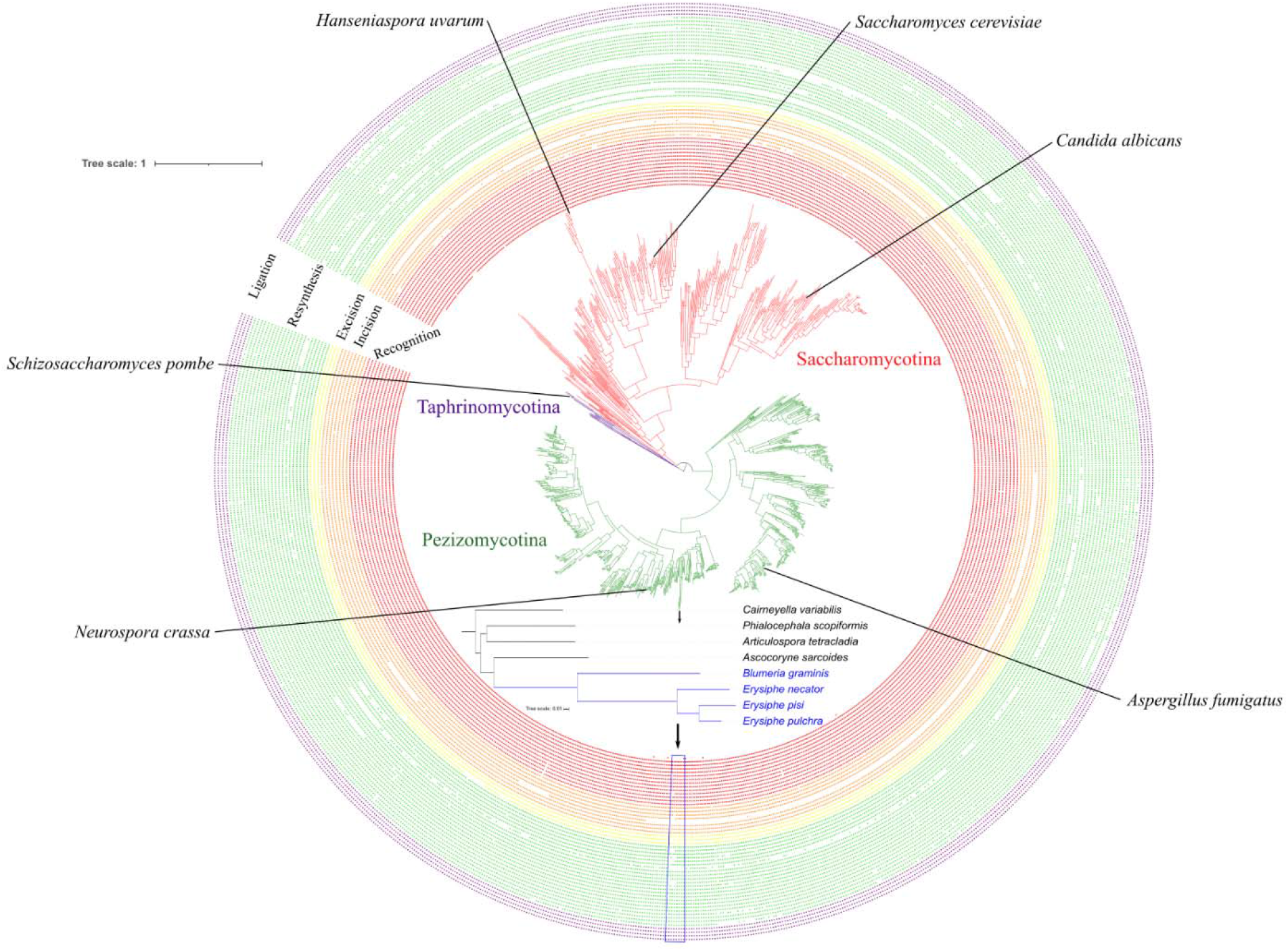
Conservation of mismatch repair (MMR) pathway genes across the fungal phylum Ascomycota. MMR genes are generally highly conserved across the phylum. A few model organisms and species of particular interest to medicine and agriculture are labeled as well as a representative species of the faster-evolving *Hanseniaspora* lineage. Gene presences are indicated in the bands surrounding the phylogeny with genes colored according to their function; red is recognition, orange is incision, yellow is excision, green is resynthesis, and purple is ligation. Branches are colored by subphylum; budding yeasts / Saccharomycotina (n = 332 species) are in red, fission yeasts / Taphrinomycotina (n = 14 species) are in purple, and filamentous fungi / Pezizomycotina (n = 761 species) are in green. Taxon names have been omitted from the phylogeny for visualization purposes; the phylogenetic tree with taxon names can be found in Figure S1 and Shen et al. (2020). The inset phylogenetic tree shows the higher loss taxa (HLT; in blue) and lower loss taxa (LLT; in black), with the blue box beneath highlighting them.

**Fig. 3:**
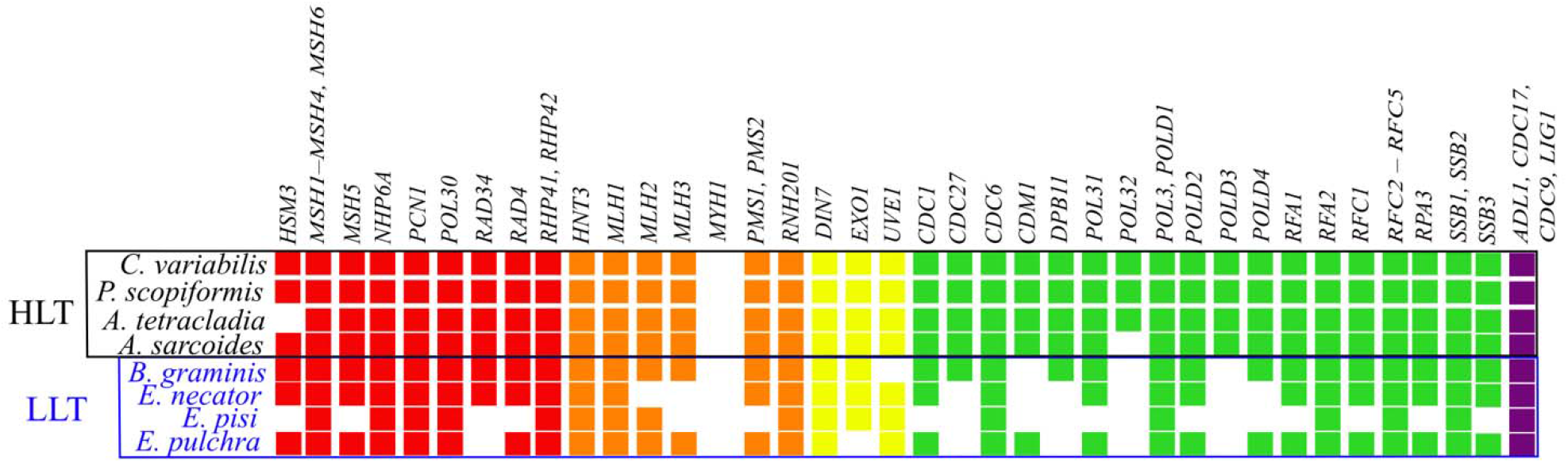
The powdery mildews *Erysiphe* and *Blumeria* have lost many more mismatch repair (MMR) pathway genes than closely related species. Higher loss taxa (HLT; shown in blue font) have lost 5 – 21 MMR genes, while lower loss taxa (LLT; shown in black font) have lost 1 – 2 genes and the median ascomycete has lost 3 genes. Note that the losses of *EXO1*, *MLH2*, *MLH3*, *MSH5*, *PMS1*, *PMS2*, and *RFC1* are uniquely observed in HLT. Genes are colored according to their function; red is recognition, orange is incision, yellow is excision, green is resynthesis, and purple is ligation.

## Results

### MMR genes are highly conserved across the fungal phylum Ascomycota

By examining the presence of 52 MMR genes using a combination of sequence similarity search algorithms across the genomes of 1,107 fungal species, we found that the MMR pathway is highly conserved across Ascomycota (a median of 49 / 52 MMR genes per species; Fig. 2; File S1). Sixteen genes were present in all species; these included five recognition genes (*MSH1*, *MSH2, MSH3, MSH4,* and *MSH6*), one incision gene (*MLH1*), one removal gene (*DIN7*), five resynthesis genes (*CDC6, RFC2, RFC3, RFC4,* and *RFC5*), and all four ligation genes (*ADL1, CDC17, CDC9*, and *LIG1*). Few genes experienced extensive loss. Of the 11 most commonly lost genes, which were lost in >5% of species, two (*MYH1* and *UVE1*) were lost in the common ancestor of Saccharomycotina, in addition to losses observed in other taxa. The remaining nine genes are unevenly distributed across functions; seven are involved in DNA resynthesis (*CDC27*, *CDM1, POL32, POLD3, POLD4, RPA3*, and *SSB3*), one is involved in mismatch recognition (*HSM3*), one is involved in incision (*HNT3*). These findings suggest that genes in the MMR pathway are well conserved across Ascomycota.

A comparison of our results with those reported in Milo et al. (2019) revealed similar patterns of gene presence and absence. For example, Milo et al. (2019) found that *MYH1* was absent from much of Pezizomycotina, which is consistent with our results. However, we did identify a few differences (inferred losses by Milo et al. (2019) vs. inferred presence in our analyses), which suggest that our pipeline is more conservative in classifying gene losses. We surmise that these differences stem from differences in the gene detection pipelines employed and the divergent objectives of the two studies; Milo et al. (2019) aimed to identify orthologs via a reciprocal best BLAST hit approach, whereas we aimed to identify homologs using pHMMs with absences verified using TBLASTN. Importantly, analysis of the human proteome using our pipeline detected all known human orthologs except for *HSM3*, *POL32*, *RPA3*, and *SSB3*, suggesting that our pipeline is well suited to detect MMR genes within Ascomycota, but that a few MMR genes may be more rapidly evolving and therefore more difficult to detect.

### Extensive loss of MMR genes in a lineage of powdery mildews

Although MMR genes are highly conserved across Ascomycota, we found that a lineage of obligate plant parasite powdery mildews have among the fewest MMR genes of the 1,107 Ascomycota species examined. We further verified gene losses in HLT and in closely related taxa that experienced fewer losses (hereafter referred to as lower loss taxa or LLT) by performing TBLASTN using the amino acid sequences found by the pHMM for each MMR gene of *Cairneyella variabilis*, an LLT with a highly conserved MMR pathway, as queries and with thresholds described by Milo et al. (2019). This resulted in 9 MMR genes losses in *Erysiphe necator*, 21 in *Erysiphe pisi*, and 7 in *Erysiphe pulchra* (Fig. 3). *E. necator* has been previously documented to have a high rates of genome evolution (Milo et al. 2019) and genomic instability (Jones et al. 2014). *Blumeria graminis*, which is sister to the *Erysiphe* genus, has lost 5 MMR genes; previous studies reported extensive gene loss in diverse pathways in this species, generally in genes thought to be unnecessary for its biotrophic lifestyle (Spanu et al. 2010). In contrast, the closely related species *C. variabilis* and *Phialocephala scopiformis* only lack *MYH1*, an adenine DNA glycosylase that is lost in most filamentous fungi (Chang et al. 2001). In addition to *MYH1*, closely related species *Articulospora tetracladia* lacks *HSM3*, which has been lost in many Pezizomycotina genomes. *Ascocoryne sarcoides* lacks *MYH1* and *POL32*, a DNA polymerase δ subunit, which is part of a larger complex that participates in multiple DNA repair pathways, including nucleotide excision repair and base excision repair (Gerik et al. 1998). Much like the rest of the phylum, genes associated with resynthesis are lost more frequently, but *Erysiphe* and *Blumeria* have lost genes associated with all MMR functions (Table S2) except ligation. In addition, seven of the observed MMR gene losses occur nowhere else in Ascomycota: *EXO1* (excision), *MLH2* (incision), *MLH3* (incision), *MSH5* (recognition), *PMS1* (incision), *PMS2* (incision), and *RFC1* (resynthesis). Taken together, these results raise the hypothesis that HLT may have a partially functional MMR pathway.

While select *Erysiphe* taxa have lost more genes than any other species, there are other species with moderate to high levels of MMR gene loss across the phylum. A total of 318 species have lost 5 or more genes across Ascomycota: 5 species in subphylum Taphrinomycotina, 239 in Saccharomycotina, and 74 in Pezizomycotina. The disproportionate number of Saccharomycotina and Taphrinomycotina species is consistent with our knowledge that organisms in these lineages have, on average, a smaller number of DNA repair genes compared to Pezizomycotina (Milo et al. 2019). MMR gene loss in certain Saccharomycotina lineages, such as in some species from the genera *Hanseniaspora* (Steenwyk, Opulente, et al. 2019)*, Tetrapisispora,* and *Dipodascus*, is comparable to the loss observed in HLT. However, only 9 other species in Pezizomycotina showed MMR gene loss to the same degree as any *Erysiphe* species. In general, species with elevated levels of gene loss primarily lost genes noted as commonly lost earlier in this paper (see “MMR genes are highly conserved across the fungal phylum Ascomycota”), with occasional losses in other genes.

There was a notable discrepancy between the presence and absence of MMR genes inferred by pHMM versus TBLASTN in HLT that was not observed in other species. When measured by pHMM, *E. pulchra* lost 42 MMR genes, as opposed to 9 when using TBLASTN with model organism sequences as queries and 7 when using TBLASTN with *C. variabilis* sequences as queries to verify the absences. *E. necator* and *E. pisi* lost 51 MMR genes according to our pHMMs, as opposed to 10 and 22 by model organism TBLASTN and 9 and 21 with *C. variabilis* TBLASTN, respectively. *B. graminis* lost 9 MMR genes by pHMM, 6 by model organism TBLASTN, and 5 by *C. variabilis* TBLASTN. In the closely related LLT *C. variabilis*, *P. scopiformis, A. tetracladia,* and *A. sarcoides,* genes deemed absent by pHMMs were also deemed absent in our model organism and *C. variabilis* TBLASTN searches; the sole exception was *HSM3*, which was identified as present in *A. sarcoides* using *C. variabilis* TBLASTN. Even though pHMMs are more sensitive in sequence similarity searches and typically outperform TBLASTN when detecting genes on an evolutionary timescale (Yoon 2009), this discrepancy is likely explained by the lower annotation quality of some HLT species and the lower genome quality for *E. pisi* (Table S1).

### Higher MMR gene loss taxa show increased number and length of microsatellites

Examination of microsatellites revealed microsatellite expansions in HLT in comparison to LLT. Specifically, we found statistically significant increases in the number and length of microsatellites in HLT compared to LLT (Fig. 4A, Tables 1, S3, and S4). Overall, after controlling for genome size, HLT had significantly more microsatellites than LLT (F = 34.83; p < 0.001; ANOVA; Table S3). This effect was driven by the very large increase in the number of mononucleotide runs in *Erysiphe* and *Blumeria* (Fig. 4C) (p < 0.001; Tukey HSD; Table S3). There was no statistically significant difference between the groups in the numbers of microsatellites with a 2-6 bp motif length (Table S3). HLT showed significantly higher average microsatellite lengths at every motif size than LLT (Fig. 4B) (p < 0.01 for 1 bp, p < 0.001 for all other motif lengths; Wilcoxon rank sum test; Table S4). HLT have an increased number of mononucleotide runs (after controlling for genome size) and an increase in length of microsatellites of all motif lengths, suggesting that the MMR pathway’s function is compromised in these species.

**Fig. 4:**
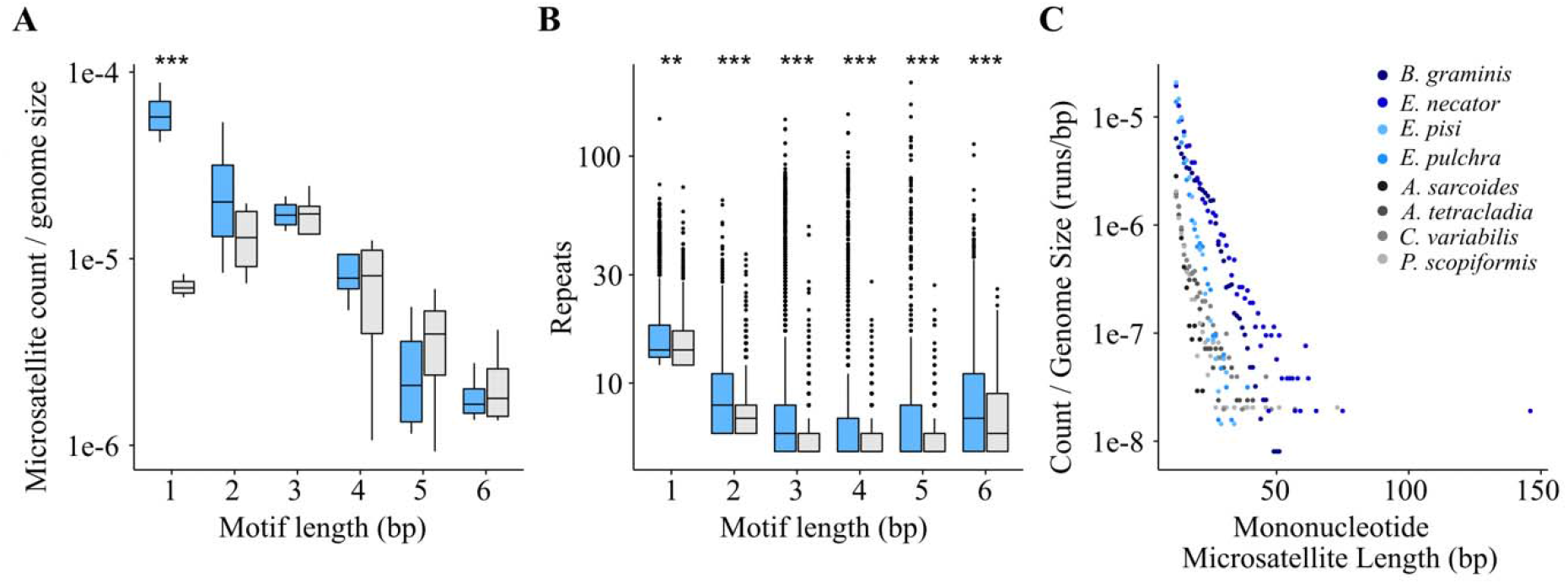
Genomes of higher loss taxa (HLT; blue bars) show a proliferation of mononucleotide runs and an increase in their microsatellite lengths compared to lower loss taxa (LLT; grey bars). (A) Examination of microsatellites in HLT and LLT (gray bars) showed a significant increase in the number of mononucleotide runs in HLT (p < 0.001; ANOVA, Tukey HSD; Table S3). Asterisks in graph indicate significance; **: p < 0.01; ***: p < 0.001. (B) Microsatellites of each motif length are significantly longer in HLT (p < 0.01 for 1 bp, p < 0.001 for all other motif lengths; Wilcoxon rank sum test; Table S4). (C) Mononucleotide runs are longer and more numerous in HLT than LLT (Tables S3 and S4).

**Table 1.**
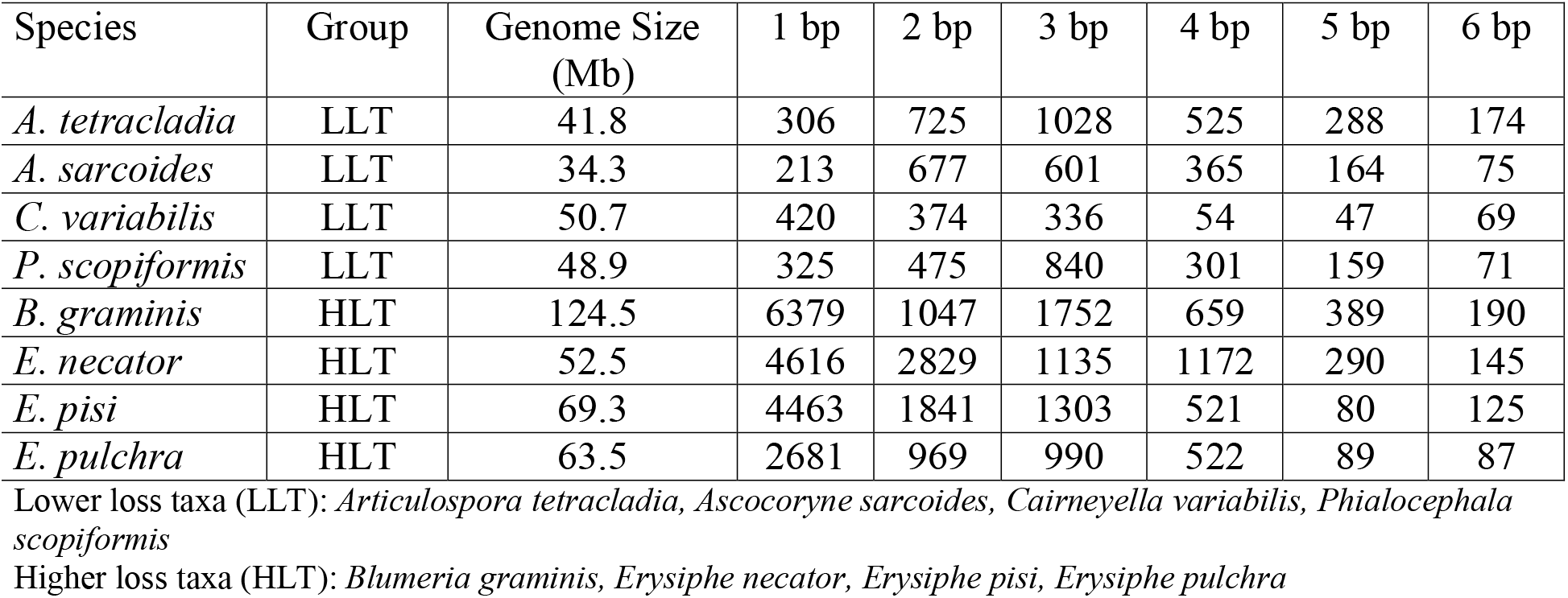
MMR gene loss is associated with an increased number of microsatellites.

### Higher loss taxa show mutational biases

By examining patterns of substitutions among HLT and LLT we found that HLT displayed stronger mutational biases associated with impaired DNA repair pathway function in comparison to LLT. For example, significantly more substitutions were observed at all codon positions in HLT vs. LLT (Fig. 5A) (p < 0.01; Tukey HSD; Table S5) and a significant bias towards substitutions in the A|T direction (Fig. 5B) (p < 0.001; Tukey HSD; Table S6). HLT also had a lower ratio of transitions to transversions (0.92 ± 0.04) than LLT (0.99 ± 0.02), though this is not statistically significant (Fig. 5C) (p = 0.06; Wilcoxon rank sum exact test; Table S7). Additionally, HLT had lower GC content (HLT: 40.10 ± 0.02% vs. LLT: 47.49 ± 0.01%). Linear regression revealed a significant decrease (F = 8.661, p = 0.026; Table S8) in GC content of the genomes as the number of MMR genes lost increases (Fig. 5D).

**Fig. 5:**
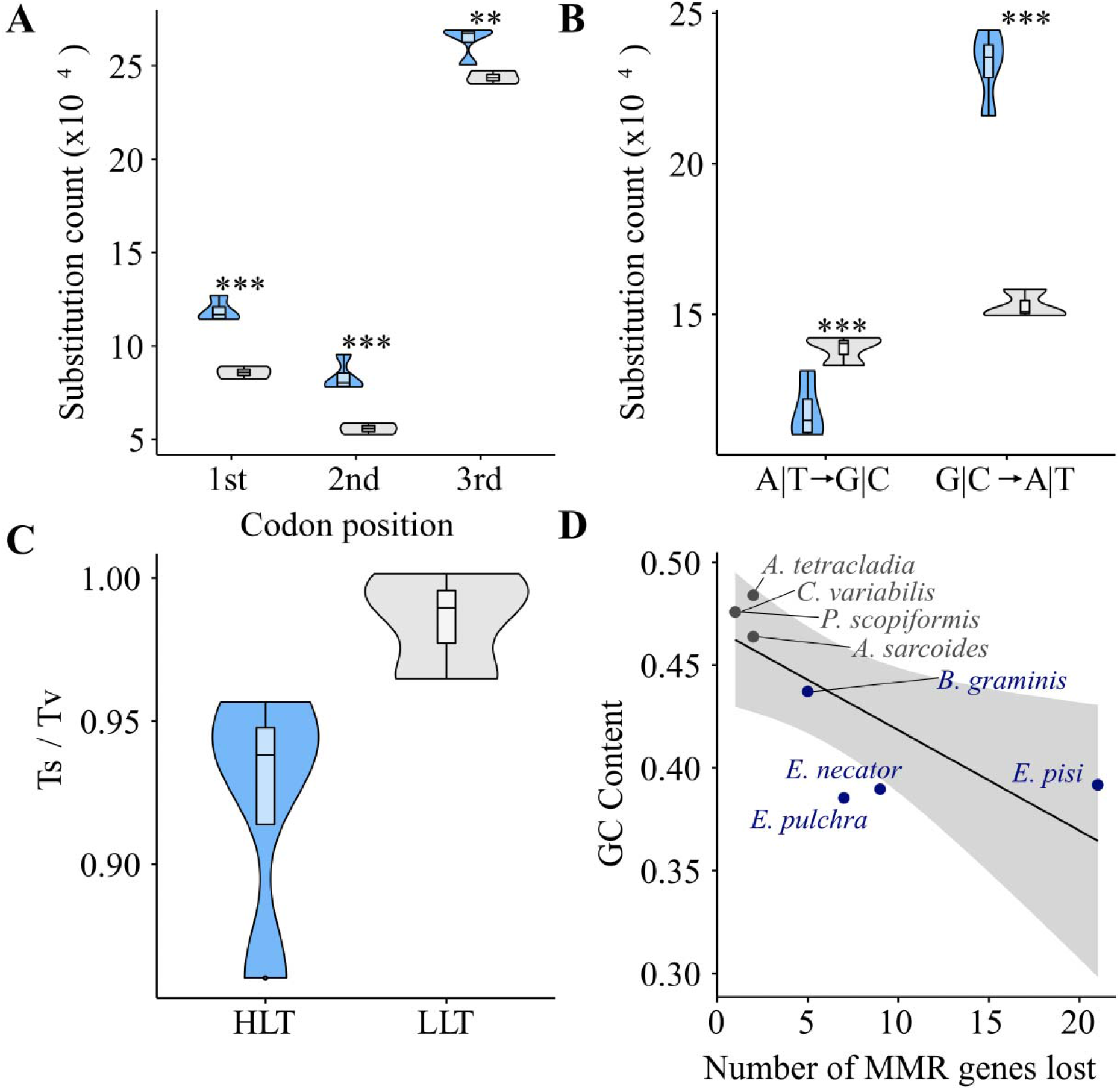
Higher loss taxa (HLT) show diverse types of mutational bias compared to lower loss taxa (LLT). (A) HLT (blue bars / fonts) show increased counts in base substitution at every codon position when compared to LLT (gray bars / fonts) (p < 0.01; ANOVA, Tukey HSD; Table S5). (B) HLT show significant mutational bias towards mutation in the A|T direction, while this trend is not significant in LLT (p < 0.001; p = 0.27; ANOVA, Tukey HSD; Table S6). (C) HLT show a decreased ratio of transitions to transversions when compared to LLT, although this difference is not statistically significant (p = 0.06; Table S7). (D) Genome GC content decreases with increasing MMR gene loss (adjusted R^2^ = 0.5225; p = 0.026; linear regression; Table S8). Asterisks in graphs indicate significance; **: p < 0.01; ***: p < 0.001.

### Higher loss taxa have experienced accelerated rates of sequence evolution

To test if the rate of evolution of HLT differed from that of LLT, we performed ω-based branch tests. Our null hypothesis was that all branches of the phylogeny for our selected eight species had the same rate of evolution, while our alternate hypothesis posited that HLT branches, including the branch of their most recent common ancestor, experienced a different rate of sequence evolution than LLT branches. We found that 60.75% of genes rejected the null hypothesis (α = 0.05; n = 500) and 39.25% failed to reject the null (n = 323) (Fig. 6A; File S2). Of the genes which rejected the null hypothesis, 79.80% (n = 399) experienced higher rates of substitution in HLT, which constitutes 48.48% of all genes tested (Fig. 6A). Among the genes that rejected the null hypothesis, the difference between the ω values for the HLT (median ω = 0.0899) and the LLT (median ω = 0.0567) showed accelerated rates of substitution for HLT branches (Fig. 6B). These results suggest that MMR gene loss is associated with a genome-wide signature of accelerated mutation rates.

**Fig. 6:**
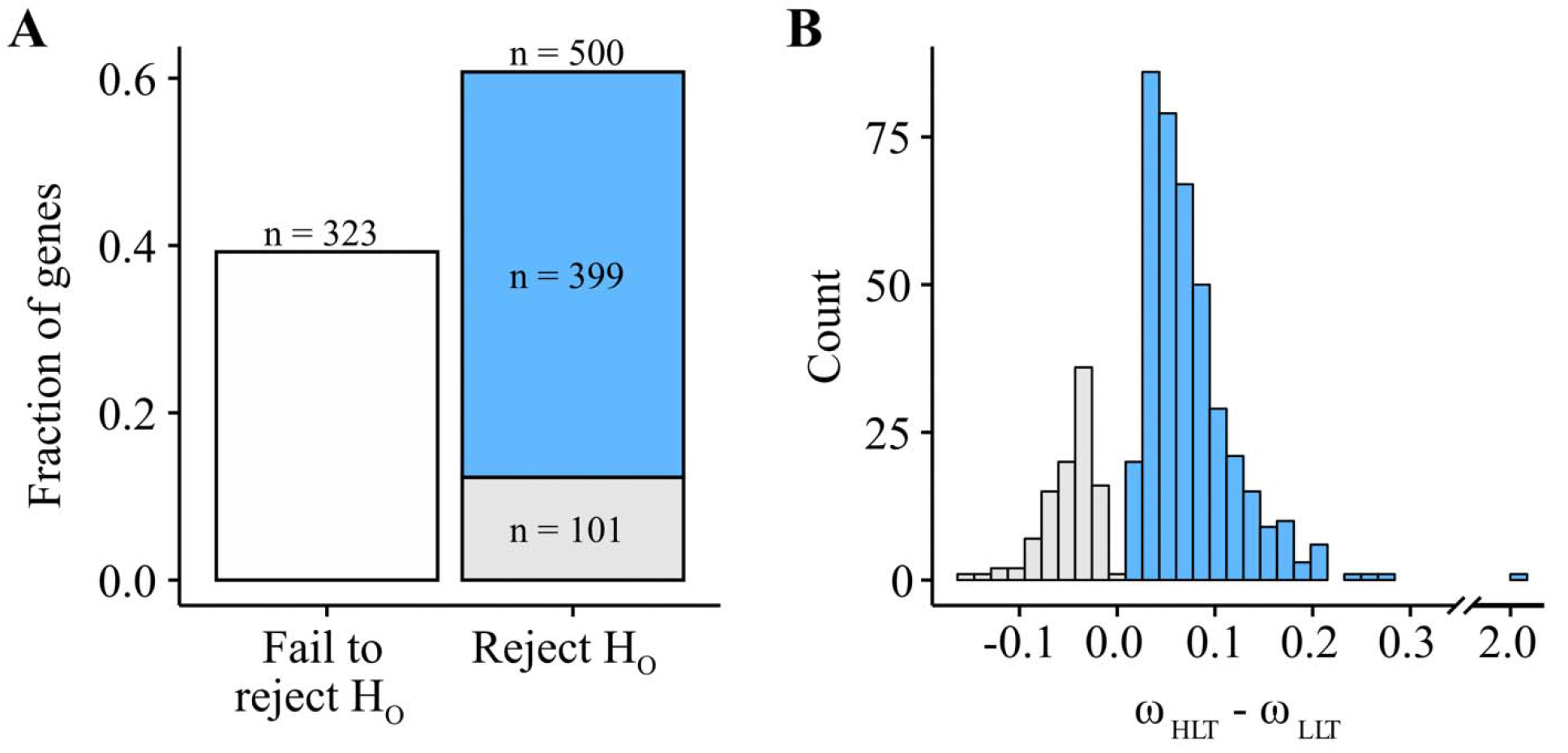
Powdery mildew higher loss taxa (HLT) show accelerated rates of evolution. (A) Most (60.75%; n = 500) BUSCO genes reject the null hypothesis that HLT branches experience the same rate of substitution as LLT branches. 48.48% of BUSCO genes support a higher rate of evolution for HLT (n = 399; in blue), 12.27% support a higher rate of evolution for LLT (n = 101; in grey), and 39.25% (n = 323) fail to reject the null hypothesis that the rate of substitution is uniform across HLT and LLT branches (in white). Among genes that support the alternative hypothesis, 79.80% (n = 399) support that *Erysiphe* and *Blumeria* evolve more quickly than LLT. (B) Among genes which reject the null hypothesis, the distribution of differences between ω values for HLT and LLT branches show elevated substitution rates in HLT.

## Discussion

Using sequence similarity searches, we examined the conservation of the MMR pathway across 1,107 ascomycete species. The near universal conservation of the vast majority of MMR genes across the phylum confirms this pathway’s known critical role for DNA maintenance (Fukui 2010; Schofield & Hsieh 2003; Kunkel & Erie 2005). However, we also discovered that a lineage of *Erysiphe* and *Blumeria* powdery mildews, named HLT, have experienced significant MMR gene loss (Fig. 3). HLT exhibit increases in mononucleotide run count, microsatellite length, mutational biases, and rate of evolution (Figs. 4, 5, and 6), suggesting that the function of their MMR pathway may be impaired. While DNA repair mechanisms are present in all eukaryotes and are highly conserved, there is mounting evidence of exceptions to this rule in the fungal kingdom (Steenwyk, Opulente, et al. 2019; Steenwyk 2021). The increased MMR gene absence observed in HLT correlates with changes in the microsatellite compositions of their genomes. The significant difference between HLT and LLT in the number of mononucleotide runs is consistent with mutational patterns present in human cancers and MMR deficient yeast cells (Arzimanoglou et al. 1998; Lang et al. 2013). Mononucleotide runs are the most prone to replication fork slippage and are used to diagnose MSI in tumors (Richman 2015). In addition to an increase in the number of mononucleotide runs (Fig. 4A), HLT showed significantly longer microsatellites for each motif length than LLT (Fig. 4B), which suggests impaired MMR function and increased replication fork slippage.

Examination of HLT genomes revealed mutational signatures suggesting that the MMR pathway has been impaired by the observed gene losses. Patterns of substitutions suggest the loss of MMR genes leads to increased substitution rates (Fig. 5A) and lower GC content (Fig. 5D). More specifically, the prominent A|T bias of substitutions in the HLT is likely driven by the known A|T bias of mutations previously observed in bacteria and eukaryotes, including *S. cerevisiae* (Hershberg & Petrov 2010; Keightley et al. 2009; Zhu et al. 2014; Lynch 2010). Furthermore, GC content decreases in proportion to the number of MMR genes lost in the HLT and LLT, which was also observed among *Hanseniaspora*, a lineage of budding yeasts that have lost diverse DNA repair genes (Steenwyk, Opulente, et al. 2019). There is no significant difference between the transition to transversion (Ts/Tv) ratios of the HLT (0.92 ± 0.04) and LLT (0.99 ± 0.02), though the trend follows what we would expect for HLT having less efficient DNA repair. Both lineages exhibit near-neutral Ts/Tv ratios (Zhu et al. 2014; Lynch et al. 2008). Examination of ω values suggests that faster rates of sequence evolution in HLT compared to the LLT may be associated with MMR gene loss. Long branches, which reflect more substitutions per site, have been previously reported elsewhere for *E. necator* (Milo et al. 2019), providing independent support to our findings.

Species in the LLT lineage show a diversity of ecologies. For example, *A. sarcoides* is saprobic fungus which grows on trees (Gianoulis et al. 2012), *A. tetracladia* is a globally-distributed aquatic hyphomycete (Seena et al. 2012), *C. variabilis* is an ericoid mycorrhizal fungus (Midgley et al. 2016), and *P. scopiformis* is a foliar endophyte (Walker et al. 2016). In contrast, species in the genera *Erysiphe* and *Blumeria* are all powdery mildews and obligate plant parasites. *B. graminis* is the only species in the genus *Blumeria* (Inuma et al. 2007), whereas there are ~450 known species in the genus *Erysiphe* (Takamatsu et al. 2015). The *Erysiphe* species we sampled are distributed across the phylogeny of the genus (Takamatsu et al. 2015; Ellingham et al. 2019); phylogenetic analyses by Takamatsu et al. (2015) placed *E. pisi* and *E. pulchra* in separate phylogenetic groups that diverged 15-20 million years ago, and analyses by Ellingham et al. (2019) showed that *E. pisi* and *E. necator* are distantly related (Ellingham et al. 2019). Considering our taxon sampling and obligate plant parasitic lifestyle of *Erysiphe* species, we hypothesize that our findings likely apply to all species in the genus.

In our approach to investigate the conservation of the MMR pathway in Ascomycota, we chose to be inclusive when selecting genes that function as part of the pathway and conservative when ruling genes absent. Some genes were computationally annotated as belonging to the MMR pathway based on sequence homology and others are implicated in multiple pathways. That said, of the 24 MMR genes lost in at least one HLT species, 16 are included in the KEGG MMR pathway entries for our three model organisms, suggesting that many of the observed losses concern genes directly involved in MMR. In addition, there may be other contributing factors to the observed mutational differences between HLT and LLT, such as dysfunction in other DNA repair pathways, loss of methyl transferases, and differences in chromatin structure (Steenwyk, Opulente, et al. 2019; Möller et al. 2021). For example, previous studies have identified the loss of genes involved in the repeat-induced point (RIP) mutation pathway in powdery mildews (Spanu et al. 2010; van Wyk et al. 2021). Loss of genes in the RIP pathway in HLT could contribute to the elevated mutation rates we observed relative to LLT. The MMR pathway is required for maintaining heterochromatin stability in *S. cerevisiae*; dysfunction in this pathway could potentially lead to genome instability within the HLT (Dahal et al. 2017). MMR is more error prone in heterochromatin than euchromatin, likely due at least in part to mechanical accessibility of the MMR machinery (Dahal et al. 2017; Sun et al. 2016). While base-base mismatches are repaired less efficiently in heterochromatin than in euchromatin, single nucleotide insertions and deletions are repaired with similar efficiency in euchromatin and heterochromatin (Dahal et al. 2017); therefore, differences in chromatin structure could have contributed to some of the observed mutational differences but not to the observed differences in mononucleotide runs and microsatellite repeats.

Loss of function in the MMR pathway may be advantageous in certain environments or under certain lifestyles. For example, strains of human pathogens with impaired MMR function are found in environments where antifungal drugs are present. Some of these strains have evolved drug resistance, so the elevated mutation rate generated by MMR gene loss may be adaptive under certain stressful situations (Billmyre et al. 2017, 2020; Rhodes et al. 2017). Species with higher levels of MMR gene loss may be associated with a parasitic lifestyle, though not all parasites have high levels of MMR gene loss; dysfunction in this pathway may be more adaptive, or at least less detrimental, to these organisms, as seen in the HLT. Loss of DNA repair pathways is also observed in other parasites and may contribute to elevated mutation rates (Gill & Fast 2007; Derilus et al. 2021). Organisms with parasitic lifestyles tend to evolve more rapidly than free-living organisms; while these mechanisms are unknown, previous work has identified that the loss of the classical nonhomologous end joining (C-NHEJ) pathway is common in this lifestyle and may even be a contributing factor (Nenarokova et al. 2019). Previous studies of genome structure in *E. necator* have found genome expansion largely driven by transposable elements and suggest that genome instability, particularly in copy number variants, can mediate rapid evolution of fungicidal resistance (Jones et al. 2014). The evolution of fungicide resistance in powdery mildews has implications for agriculture; major crops are impacted by these pathogens and some are able to quickly evolve resistance to antifungal chemicals, with resistance evolution accelerated by increased use (Jones et al. 2014; Vielba-Fernández et al. 2020). More broadly, genome instability among HLT taxa reflects their parasitic lifestyle, which is associated with gene loss and plastic genomic architecture (Schmidt & Panstruga 2011). Gene loss in primary and secondary metabolism, enzymes acting on carbohydrates, and transporters has been documented in *B. graminis*, as well as massive expansion in retrotransposons and genome size, reflecting extreme genomic changes associated with its parasitic lifestyle (Spanu et al. 2010). The lost genes are involved in diverse pathways, including anaerobic fermentation, biosynthesis of glycerol from glycolytic intermediates, and nitrate assimilation, and include multiple subfamilies of transporters (Spanu et al. 2010). Given their extreme genomic changes and importance to agriculture, *Blumeria* and *Erysiphe* may be novel models to study the outcome and evolutionary trajectory of sustained loss of MMR pathways.

## Methods

### Curation of the set of DNA mismatch repair pathway genes

To investigate the presence and absence of MMR genes across the fungal phylum Ascomycota, we curated a dataset of MMR genes from the genomes of three fungal model organisms representing the three different subphyla: *Saccharomyces cerevisiae* (subphylum Saccharomycotina), *Neurospora crassa* (Pezizomycotina), and *Schizosaccharomyces pombe* (Taphrinomycotina). We used three sources to curate genes that are part of the MMR pathway: the Kyoto Encyclopedia of Genes and Genomes (KEGG, genome.jp/kegg/; Kanehisa & Goto, 2000), the *Schizosaccharomyces pombe* database (PomBase, pombase.org/; Lock et al., 2019; The Gene Ontology Consortium, 2019), and the *Saccharomyces* Genome Database (SGD, yeastgenome.org/; Cherry et al., 2012). Aiming to be inclusive when selecting genes to be included as MMR, genes in the KEGG diagram of the MMR pathway for each species were included and the gene ontology (GO) term “mismatch repair” was used to search for the genes on SGD and Pombase (Ashburner et al. 2000). We used both computationally and manually curated genes from SGD. We began curating our set of MMR genes in *S. cerevisiae*, with a total of 30 MMR genes identified with KEGG and SGD. Next, we searched KEGG and Pombase for genes in *S. pombe* that had not been annotated as part of the MMR pathway in *S. cerevisiae* (n = 15). We concluded by searching for *N. crassa* MMR genes in KEGG which had not already been categorized as MMR genes in the other two species (n =7). KEGG listed two sequences for the *N. crassa* gene *LIG1*; however, since our sequence similarity search analyses with both sequences yielded identical patterns of loss, we present them as one gene. This approach yielded a total of 52 genes associated with MMR (Table S2).

### MMR gene conservation analysis

To examine the conservation of MMR genes across Ascomycota, we implemented a sensitive probabilistic modeling approach using profile Hidden Markov Models (pHMMs) (Johnson et al. 2010) of MMR genes across the genomes of 1,107 species (Shen et al. 2020). To construct pHMMs, we first searched for putative homologs of MMR genes in the fungal RefSeq protein database using the blastp function of BLAST+, v2.8.1, with a bitscore threshold of 50 and an e-value cutoff of 1 × 10^−3^ (Pearson 2013). We retrieved the top 100 hits using SAMtools, V1.6 (Li et al. 2009) with the ‘faidx’ function. We used MAFFT, v7.402 (Katoh et al. 2002), with the ‘genafpair’ and ‘reordered’ parameters, a maximum of 1000 cycles of iterative refinement, the BLOSUM62 matrix, a gap opening penalty of 1.0, and the retree parameter set to 1, to align the sequences following previously established protocol (Steenwyk, Shen, et al. 2019).We then used the aligned amino acid sequences as input to the ‘hmmbuild’ function in HMMER-3.1B2 to construct each pHMM. We ran the pHMMs of the 52 proteins against all 1,107 proteomes using the ‘hmmsearch’ function. For a gene to be considered present, we set a bitscore threshold of at least 50 and an e-value threshold of less than 1 × 10^−10^. We used the TBLASTN function of BLAST+, V2.8.1 with a bitscore threshold of 50, e-value cutoff of 1 × 10^−6^, and 50% minimum query coverage to verify MMR gene absence using the protein sequence of the gene in question and the 1,107 Ascomycota genomes. We used the Interactive Tree of Life (iTOL), v4 (Letunic & Bork 2019) to visualize the conservation of MMR genes on the Ascomycota phylogeny and to map losses on it. To further verify gene absences in HLT and LLT with *C. variabilis* amino acid sequences, we used the TBLASTN function of BLAST+, V2.8.1 with an e-value threshold of less than 1 × 10^−5^, a word size of 5 or more, and minimum query coverage of 80% (Milo et al. 2019).

### Microsatellite identification and characterization

To identify microsatellites and evaluate their numbers and lengths between genomes with substantial MMR gene loss against those with higher levels of MMR gene conservation, we used the Microsatellite Identification tool (MISA), v2.0 (Beier et al. 2017). Specifically, we compared the microsatellites of two groups of taxa, each of which contained four species. The group of higher loss taxa (HLT) contains the powdery mildews *Blumeria graminis, Erysiphe necator, Erysiphe pisi*, and *Erysiphe pulchra*, which show high levels of MMR gene loss relative to other ascomycetes. The group of lower loss taxa (LLT) contains four closely related species with low levels of MMR gene loss, similar to patterns seen across the rest of the phylum: *Articulospora tetracladia, Ascocoryne sarcoides, Cairneyella variabilis,* and *Phialocephala scopiformis*. The length minimums used for MISA to identify a microsatellite are as follows: 1 base pair (bp) motifs must repeat 12 times, 2 bp motifs must repeat 6 times, 3-6 bp motifs must repeat 5 times. All values used are MISA defaults, except the mononucleotide parameter, which was increased from the default value of 10 repeats to 12 (Beier et al. 2017; Temnykh et al. 2001). A 2-way ANOVA test was performed to test for significance in the number of microsatellites controlled by genome size of each motif length between HLT and LLT. If the 2-way ANOVA rejected the null hypothesis (α = 0.05), pairwise comparisons were made with the Tukey Honest Significant Differences (HSD) test. We performed the statistical analysis using R, v3.4.1 (https://www.r-project.org/) and made the figures using ggplot2, v3.1.0 (Wickerham 2016), and ggpubfigs, v1.0.0 (Steenwyk 2020).

### Estimation of mutational bias and rate of sequence evolution

To characterize the mutational spectra and estimate the rate of sequence evolution between HLT and LLT, we first identified and aligned orthologous sequences across all eight genomes. Orthologous single-copy protein sequences from genes present in all eight genomes (n = 823) were identified using the BUSCO, v4.0.4 (Waterhouse et al. 2018) pipeline and the OrthoDB, v10, Ascomycota database (Creation date: 2019-11-20) (Kriventseva et al. 2019). We hereafter refer to the 823 single-copy genes as BUSCO genes. BUSCO genes were aligned using MAFFT, v7.402 (Katoh et al. 2002), using the same settings described above. Codon-based alignments were generated by threading the corresponding nucleotide sequences onto the protein alignment using ‘thread_dna’ function in PhyKIT, v0.1 (Steenwyk et al. 2020).

To examine patterns of substitutions, we used codon-based alignments to identify nucleotides that differed in a given taxon of interest compared to *C. variabilis*, which was the sister taxon to a clade comprised of the other seven genomes of interest in the Ascomycota phylogeny. More specifically, we compared the character states for a species of interest to *C. variabilis* for each site of each alignment, tracking codon position information (i.e., first, second, or third codon position). We also determined if the substitution was a transition or transversion and examined substitution directionality (e.g., A|T to G|C or G|C to A|T) using *C. variabilis* as the outgroup. These analyses were completed using custom python scripts that utilize functions in Biopython, v1.70 (Cock et al. 2009).

Finally, we used the codon alignments to compare the rate of sequence evolution between HLT and LLT. Specifically, we measured the ratio of the rate of nonsynonymous substitutions to the rate of synonymous substitutions (dN/dS or ω) along the species phylogeny for each gene using the CODEML function in PAML, v4.9 (Yang 2007). For each test, the null hypothesis (H_o_) was that all branches had the same ω value (model = 0); the alternative hypothesis (H_A_) was that all HLT branches, including the branch of their most recent common ancestor, had one ω value and all other branches had a distinct ω value (model = 2). To determine if the alternative hypothesis was a better fit than the null hypothesis (α = 0.05) we used a likelihood ratio test.

## Acknowledgements

We thank Qianhui Zheng for performing exploratory analysis on the HLT and LLT genomes.

## Data availability statement

Supporting statistical analysis, the Ascomycota phylogeny with species names, and 2 supplementary data files (MMR gene presence/absence matrix and ω output) are available via figshare at https://doi.org/10.6084/m9.figshare.14410994. The data supporting the phylogeny of Ascomycota are available at https://doi.org/10.6084/m9.figshare.12751736.

